# Morphological brain networks of white matter: mapping, evaluation, characterization and application

**DOI:** 10.1101/2023.06.02.543370

**Authors:** Junle Li, Zhen Li, Yuping Yang, Zhenzhen Luo, Yaou Liu, Jinhui Wang

## Abstract

Neuroimaging-based connectomics studies have long focused on the wiring patterns between gray matter regions. In recent years, increasing evidence emerges that neural activity in specific sets of white matter (WM) tracts dynamically fluctuates in a coordinated manner. However, the structural basis underlying the coordination is poorly understood largely due to the lack of approaches for estimating structural relations between WM regions. Here, we developed an approach to construct morphological WM networks based on structural magnetic resonance imaging. We found that the morphological WM networks exhibited nontrivial organizational principles, presented good to excellent short- and long-term reliability, accounted for phenotypic interindividual differences (Motor and Cognition), and were under genetic control. Interestingly, highly heritable edges contributed largely to interindividual differences in phenotype. Through integration with other multimodal and multiscale data, we further showed that the morphological WM networks were able to predict regional profiles of hamodynamic coherence, metabolic synchronization, gene co-expression and chemoarchitectonic covariance. Moreover, the prediction followed functional connectomic hierarchy of WM for hamodynamic coherence, was driven by genes enriched in the forebrain neuron development and differentiation for gene co-expression, and was attributed to serotonergic system-related receptors and transporters for chemoarchitectonic covariance. Finally, applying our approach to multiple sclerosis and neuromyelitis optica spectrum disorders, we found that both diseases were associated with morphological WM dysconnectivity, which was correlated with clinical variables and able to diagnose and differentiate the diseases. Altogether, our findings indicate that morphological WM networks provide a reliable and meaningful means to explore WM architecture in health and disease.

## Introduction

The human brain operates essentially as an interconnected complex network in favor of behavior and cognition (1). This feature makes the brain particularly amenable to research from the perspective of modern network theory (2). To date, great progress has been made in the last decade in uncovering organizational principles that govern the human brain networks and adaptive alterations of the networks in development, aging and diseases (3–8). However, current attention of human brain network studies is mainly focused on the wiring patterns between gray matter regions, which are typically thought to be responsible for brain function.

In addition to gray matter, white matter (WM), accounting for nearly half of adult brain, is another important type of brain tissue composed of neuronal axons coated with electrical insulation called myelin. WM tracts play crucial roles in conveying sensory and motor information, mediating interhemispheric communication, and connecting various cortical areas. Thus, it is not surprising to find that neural activities as measured by blood oxygen level-dependent functional magnetic resonance imaging (BOLD-fMRI) are strongly correlated between segmented WM tracts and specific cortical regions (9). Intriguingly, besides the functional interactions between WM and gray matter, recent studies demonstrate that neural activities in WM themselves are also organized as large-scale functional networks with synchronous signal fluctuations in specific sets of WM tracts (10–12). Moreover, the functional WM networks are test-retest (TRT) reliable, exhibit non-random topology, relate to phenotypic differences between individuals, and are altered in brain disorders (12–16). These findings collectively indicate that neural activities are encoded in WM circuits, and a comprehensive characterization of functional WM networks is important for fully understanding the whole-brain organization (17). However, although functional WM networks are increasingly studied, very little is known regarding their structural basis largely due to the lack of approaches for estimating structural connectivity between WM regions.

Straightforwardly, structural connectivity can be obtained via diffusion-weighted tractography to reconstruct axonal tracts between WM regions. However, whole-brain tractography remains challenging as embodied in underestimation of long-distance projections (18), numerous false-positive connections (19), susceptibility to noise (e.g., head motion) during image acquisition (20). Recently, evidence from both molecular biology and in vivo neuroimaging studies indicates that neuronal or regional morphology is informative to infer their structural connectivity (21, 22). Moreover, similarity in microcosmic cytoarchitecture is directly correlated with cortico-cortical structural connectivity derived from macroscomic neuroimaging data (23). Based on these findings, we hypothesize that axonal morphology can be used to indirectly study structural connectivity between WM regions. The morphology-informed analysis of structural connectivity may be particularly appropriate for WM regions given their relatively homogeneous cellular composition (oligodendrocytes and astrocytes).

In this study, we proposed an approach to construct WM networks by mapping interregional similarity based on regional morphology (i.e., WM volume and deformation) derived from structural MRI data. For the constructed networks (hereafter referred to as morphological WM networks), we first depicted their topological organization using graph-based complex network measures. Then, we evaluated both short-term and long-term TRT reliability of the morphological WM networks. Afterwards, we examined phenotypic associations of the morphological WM networks with a broad range of behavioral and cognitive measures, and evaluated the extent to which the morphological WM networks were under genetic control. After these analyses, the morphological-functional relationship of WM networks was explored by using a set of communicational organization and morphological similarity profiles to separately predict regional hamodynamic coherence profile derived from resting-state BOLD-fMRI (R-BOLD-fMRI) data and metabolic synchronization profile derived from resting-state 18F-fluorodeoxyglucose positron emission tomography (R-FDG-PET) data. To better understand the morphological WM networks, we further linked them with brain-wide transcriptional profiles and neurotransmitter receptor and transporter distributions to analyze their genetic and chemoarchitectonic correlates. Finally, we applied the morphological WM networks to a multicentre dataset of multiple sclerosis (MS) and neuromyelitis optica spectrum disorders (NMOSD) to examine their clinical value in helping diagnose and differentiate the two diseases. Based on our findings, we argue that morphological WM networks provide a TRT reliable, phenotypically related, genetically originated, functionally relevant, neurobiologically meaningful and clinically valuable approach for future studies of human brain WM in both health and disease.

## Results

### General description of datasets, analytical methods, and research questions

In this study, we proposed an approach to construct morphological WM networks using structural MRI data. Briefly, for a structural MRI image, a voxel-based morphometry method was first used to derive a WM volume map and deformation map. These two maps were then separately used to construct a morphological WM network (i.e., volume-based network, VBN, and deformation-based network, DBN) by estimating interregional similarity among 48 WM regions of interest (ROIs) (24) via a Jensen-Shannon divergence-based similarity (JSDs) measure (25). For the constructed morphological WM networks, seven independent datasets were subsequently utilized to depict, evaluate and characterize them from different perspectives, involving organizational principle, TRT reliability, phenotypic association, heritability, functional relevance, neurobiological substrate, and clinical diagnostic value. Specifically, the Human Connectome Project (HCP) dataset (www.humanconnectome.org) (26) was used to investigate the topological organization, phenotypic association, heritability and relationship with hamodynamic coherence networks of morphological WM networks; the Hangzhou Normal University (HNU) dataset (http://dx.doi.org/10.15387/fcp_indi.corr.hnu1) (27) and South West University (SWU) dataset (http://dx.doi.org/10.15387/fcp_indi.retro.slim) (28) were used to examine the short-term and long-term TRT reliability of morphological WM networks, respectively; the Monash University (MU) dataset (https://openneuro.org/datasets/ds002898/versions/1.1.0) (29) was used to explore the association of morphological WM networks with metabolic synchronization networks; the JuSpace dataset (https://github.com/juryxy/JuSpace) (30) was used to construct a chemoarchitectonic WM network for analyzing molecular relevance of morphological WM networks; the Allen Human Brain Atlas (AHBA) dataset (http://human.brain-map.org/) (31) was used to construct a transcriptional WM network for examining genetic correlate of morphological WM networks; and the MS and NMOSD multicentric dataset (32) was used to study clinical diagnostic value of morphological WM networks.

### Similarity patterns of morphological WM networks

We constructed the morphological WM networks based on the 444 unrelated healthy participants in HCP dataset, and showed the similarity patterns.

Fig 1A shows the group-level mean morphological similarity matrices of WM. In general, the interregional morphological similarities were high for most connections regardless of the choice of morphological index and whether spatial smoothing was performed. Nonetheless, specifically organized patterns were obvious. In particular, the mean interregional morphological similarity for homotopic connections linking geometrically corresponding regions between the two hemispheres was consistently found to be significantly higher than that for heterotopic connections linking non-homotopic regions (*T* test; spatial smoothing: *T_859_* = 3.257, *P* = 0.001 for the VBN and *T_859_* = 3.364, *P* < 0.001 for the DBN; no spatial smoothing: *T_859_* = 1.984, *P* = 0.048 for the VBN and *T_859_*= 1.784, *P* = 0.075 for the DBN).

**Fig 1.**
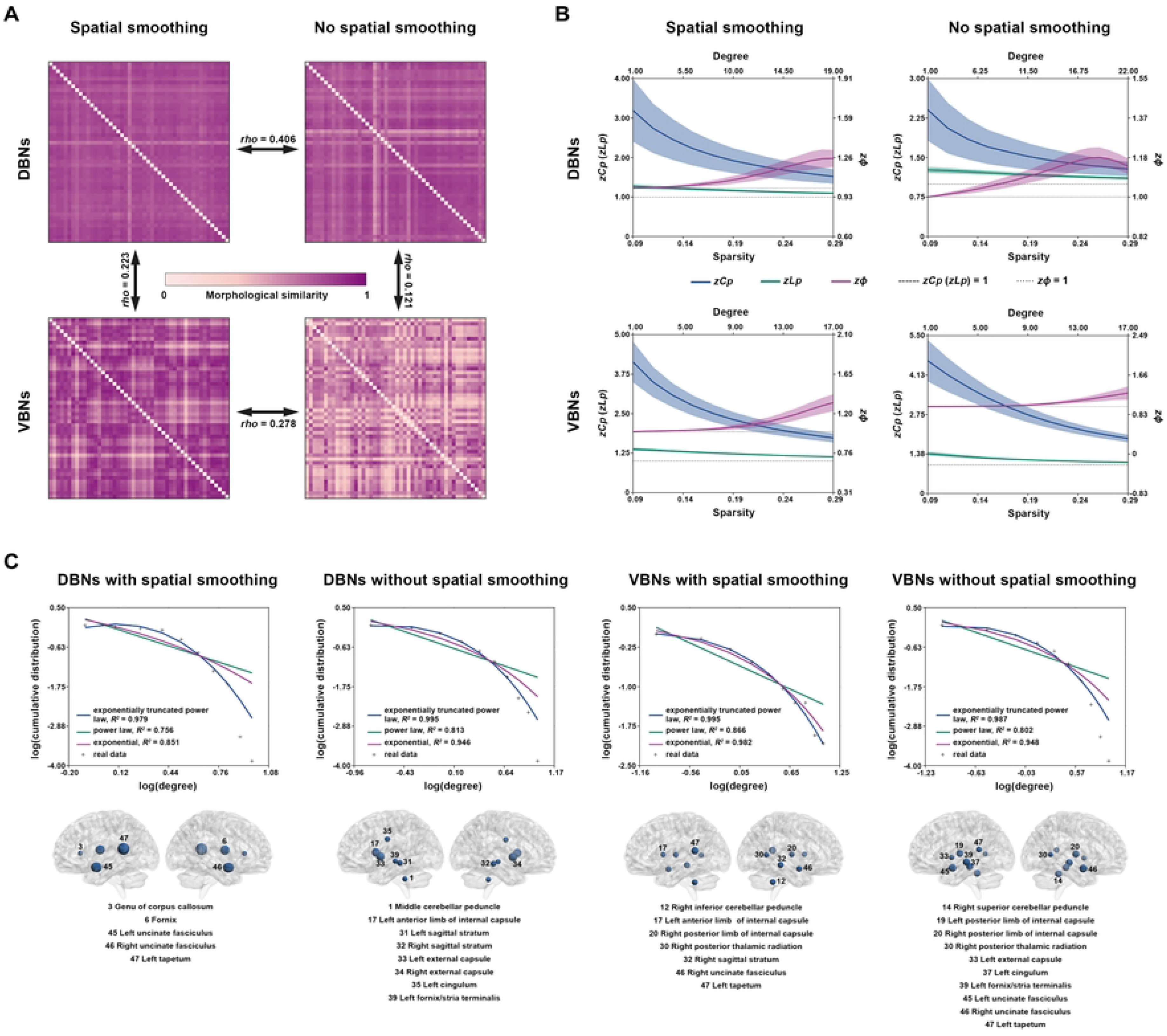
Topological descriptions of morphological WM networks. **A**) Mean interregional morphological similarity matrices and their cross-analytical strategy spatial similarities. **B**) Global organization of morphological WM networks. Compared with matched random networks, the morphological WM networks exhibited higher clustering coefficient and approximately equal characteristic path length, and higher rich-club coefficient, indicative of small-world organization and rich-club architecture. **C**) Local organization of morphological WM networks. The degree distribution of the group-level mean morphological WM networks was best fitted by the exponentially truncated power law model (upper panel), indicative of the existence of highly connected hubs (lower panel). Notably, the right uncinated fasciculus and left tapetum were frequently identified as hubs. DBNs, deformation-based networks; VBNs, volume-based networks; *zC_p_*, normalized clustering coefficient; *zL_p_*, normalized characteristic path length; *zΦ*, normalized rich-club coefficient.

We further examined the similarities and differences between the group-level mean morphological similarity matrices. Significant but relatively low Spearman rank correlations were observed between the VBN and DBN (spatial smoothing: *rho* = 0.223, *P* < 0.001; no spatial smoothing: *rho* = 0.121, *P* = 0.003) and between spatially smoothed and unsmoothed data (VBN: *rho* = 0.278, *P* < 0.001; DBN: *rho* = 0.406, *P* < 0.001). Two-way repeated ANOVA revealed that morphological index (*F_1,1127_*= 1118.672, *P* < 0.001) and spatial smoothing (*F_1,1127_*= 1773.517, *P* < 0.001) significantly affected interregional morphological similarity of the WM networks in an interactive manner (*F_1,1127_* = 1185.991, *P* < 0.001). Post hoc comparisons revealed that the DBN had significantly higher morphological similarity than the VBN no matter whether spatial smoothing was performed (spatial smoothing: *T_1127_* = 5.195, *P* < 0.001; no spatial smoothing: *T_1127_* = 42.597, *P* < 0.001) and spatial smoothing was associated with significantly higher morphological similarity for both the VBN (*T_1127_* = 40.970, *P* < 0.001) and DBN (*T_1127_* = 12.136, *P* < 0.001). These findings suggest distinct wiring patterns of the morphological WM networks between the VBN and DBN and profound effects of spatial smoothing on the morphological WM networks.

### Topological organization of morphological WM networks

We constructed the morphological WM networks based on the 444 unrelated healthy participants in HCP dataset, and explored the topological organization.

#### Small-world organization

Each morphological WM network exhibited typical small-world behaviors regardless of the choice of morphological index and whether spatial smoothing was performed, characterized by the combination of normalized clustering coefficient > 1 and normalized characteristic path ∼ 1 (Fig 1B).

#### Rich-club architecture

Each morphological WM network exhibited a normalized rich-club coefficient > 1 over a consecutive range of degree thresholds regardless of the choice of morphological index and whether spatial smoothing was performed, a typical feature of rich-club organization (Fig 1B; sparsity = 0.27).

#### Hubs

The exponentially truncated power law model gave the best fitting to the degree distribution of the group-level mean morphological WM networks regardless of the choice of morphological index and whether spatial smoothing was performed (Fig 1C). These findings indicate the existence of highly connected hubs in the morphological WM networks (Fig 1C). Of note, the right uncinated fasciculus and left tapetum were frequently identified as hubs.

### TRT reliability of morphological WM networks

We calculated the intraclass correlation coefficient (*ICC*) value of morphological similarity, based on the HNU dataset with the each scan every three days over one month and SWU dataset with the average time intervals as 304.14, 515, and 817.87 days between the first and second scans, second and third scans, and first and third scans, respectively, for short-term and long-term TRT reliability.

#### High TRT reliabilities of morphological WM networks

Good to excellent TRT reliabilities were observed for most connections in the morphological WM networks irrespective of the choice of morphological index, the implementation of spatial smoothing and time interval of scan-rescan (Fig 2A). For example, the mean (standard deviation) ICC values of the VBNs were 0.832 (0.115) and 0.844 (0.103) for short-term and long-term time intervals, respectively, when spatial smoothing was performed.

**Fig 2.**
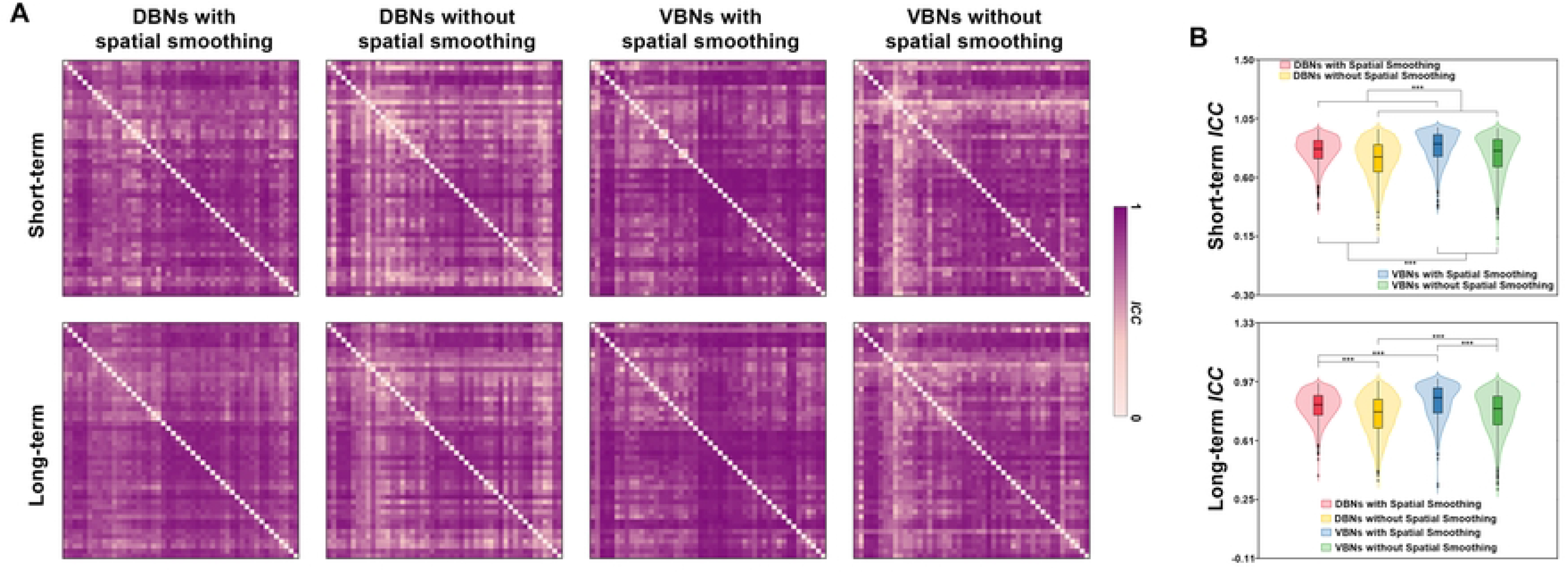
TRT reliabilities of morphological WM networks. **A**) Good to excellent short-term and long-term TRT reliabilities were observed for most connections in the morphological WM networks. **B**) Violin plots showing effects of different analytical strategies on the TRT reliabilities of the morphological WM networks. The implementation of spatial smoothing significantly increased the reliabilities of the WM networks, and the VBNs exhibited significantly higher reliabilities than the DBNs. ***, *P* < 0.001; DBNs, deformation-based networks; VBNs, volume-based networks; *ICC*, intraclass correlation coefficient.

#### Effects of different analytical strategies on the TRT reliabilities of morphological WM networks

Two-way repeated ANOVA revealed significant main effects of morphological index (*F_1,1127_* = 120.381, *P* < 0.001) and spatial smoothing (*F_1,1127_* = 610.157, *P* < 0.001) on the short-term TRT reliabilities of the WM networks. Post hoc comparisons revealed that the VBNs exhibited significantly higher short-term reliabilities than the DBNs (*T_2255_* = 12.182, *P* < 0.001), and the implementation of spatial smoothing significantly increased the short-term reliabilities of the WM networks (*T_2255_* = 26.276, *P* < 0.001). For long-term TRT reliabilities, morphological index (*F_1,1127_* = 91.805, *P* < 0.001) and spatial smoothing (*F_1,1127_* = 722.086, *P* < 0.001) significantly affected the WM networks in an interactive manner (*F_1,1127_*= 8.626, *P* = 0.003). Post hoc comparisons revealed that the VBNs exhibited significantly higher long-term reliabilities than the DBNs no matter whether spatial smoothing was performed (spatial smoothing: *T_1127_*= 10.442, *P* < 0.001; no spatial smoothing: *T_1127_* = 5.829, *P* < 0.001), and the implementation of spatial smoothing significantly increased the long-term reliabilities of both the VBNs (*T_1127_* = 21.702, *P* < 0.001) and DBNs (*T_1127_* = 20.991, *P* < 0.001) (Fig 2B).

Based on these findings, only the WM networks (VBNs and DBNs) derived from spatially smoothed data were used for the following analyses.

### Behavioral and cognitive correlation and prediction

Based on the constructed morphological WM networks and organized behavioral and cognitive domains (Alertness, Cognition, Emotion, Motor, Personality, and Sensory), we examined the ability of morphological WM networks to explain interindividual variance and predict individual performance in specific behavioral and cognitive domains via the partial least-squares (PLS) regression and brain bias set (BBS) modeling method (33).

#### Morphological WM networks can explain interindividual variance in specific behavioral and cognitive domains

The VBNs explained significant proportions of interindividual variance in the Cognition (15.1%, *P* = 0.006) and Motor (15.1%, *P* = 0.006) domains (Fig 3 left). No significant results were found for the DBNs.

**Fig 3.**
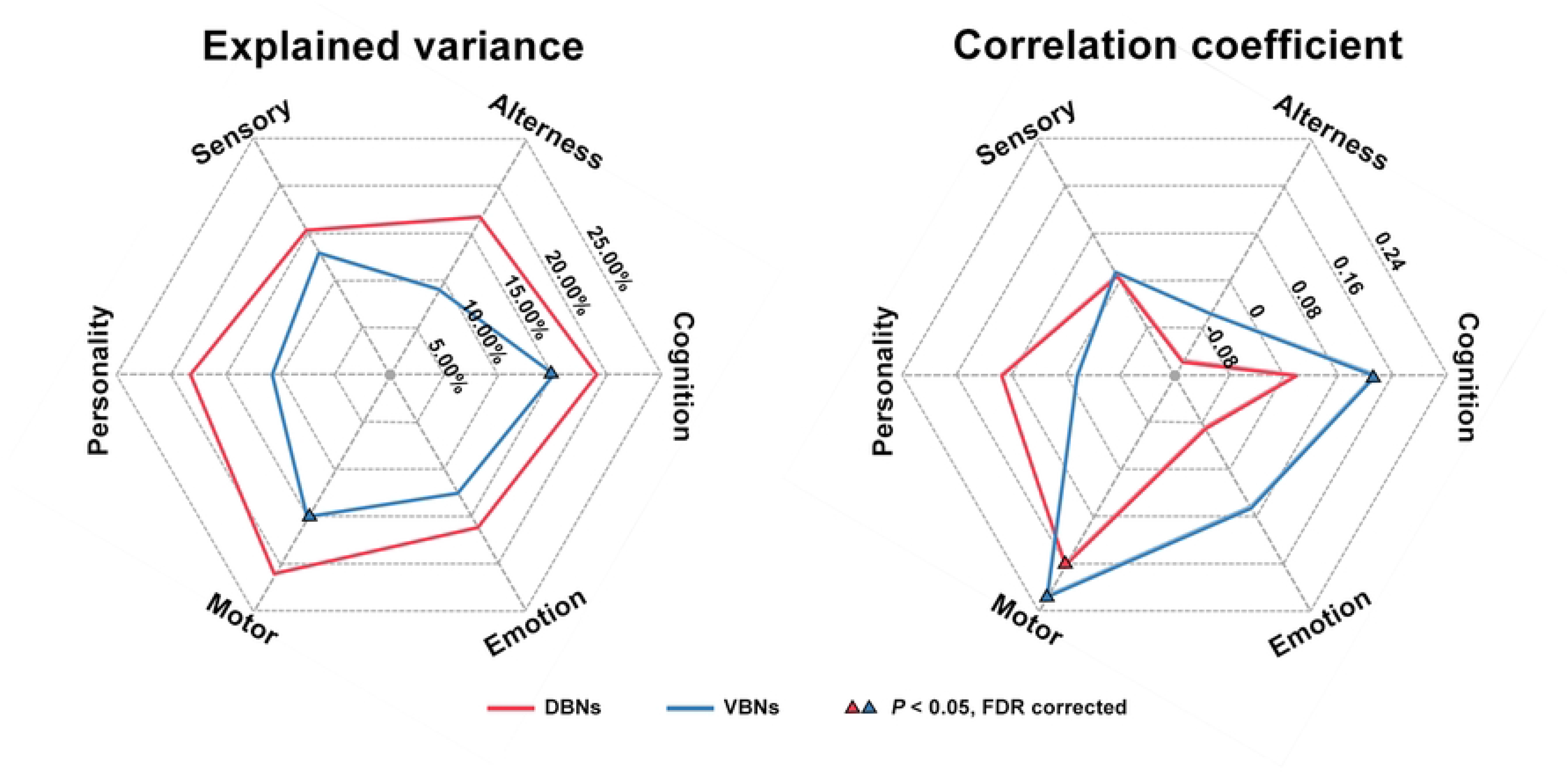
Behavioral and cognitive associations of morphological WM networks. The morphological WM networks explained significant proportions of interindividual variance in the Motor and Cognition domains (left) and predicted individual performance in these two domains (right). DBNs, deformation-based networks; VBNs, volume-based networks; FDR, false discovery rate.

#### Morphological WM networks can predict individual performance in specific behavioral and cognitive domains

The VBNs predicted individual scores of the Cognition (mean *r* = 0.134, *P* = 0.013) and Motor (mean *r* = 0.215, *P* < 0.001) domains, while the DBNs predicted individual scores of the Motor domain (mean *r* = 0.161, *P* = 0.005) (Fig 3 right).

### Heritability of morphological WM networks

A genetic ACE model was used to investigate the extent to which morphological WM networks were genetically controlled, based on the 217 pairs of monozygotic (MZ) and dizygotic (DZ) twins in HCP dataset.

Morphological WM networks exhibited overall low-moderate heritability for both the VBNs (0.382 ± 0.195) and DBNs (0.207 ± 0.139). Compared with the DBNs, the VBNs were associated with significantly higher heritability (*T_1127_* = 29.337, *P* < 0.001). Moreover, the heritability showed spatially heterogeneous distributions with a specific set of edges under strong genetic control (VBNs: 586 edges, 0.522 ± 0.125; DBNs: 137 edges, 0.359 ± 0.112) [*P* < 0.05, false discovery rate (FDR) corrected] (Fig 4). Interestingly, we found significantly positive Spearman correlations for the VBNs between the edge’s heritability and contribution to explaining interindividual variance in the Motor (*rho* = 0.273, *P* < 0.001) and Cognition (*rho* = 0.126, *P* = 0.024) domains, and predicting individual performance in the Cognition domain (*rho* = 0.109, *P* = 0.021).

**Fig 4.**
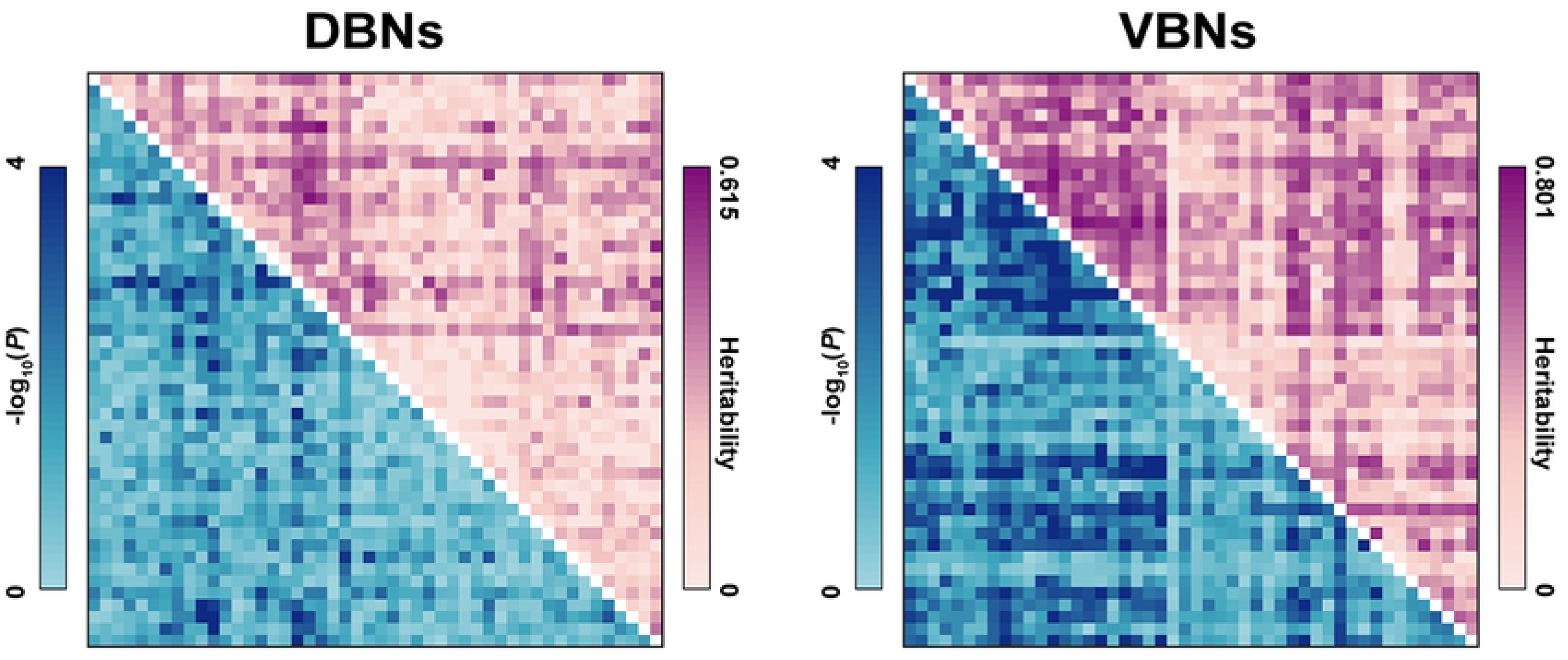
Heritability of morphological WM networks. Morphological WM networks exhibited overall low-moderate heritability with a specific set of edges under strong genetic control. Moreover, the VBNs exhibited significantly higher heritability than the DBNs. DBNs, deformation-based networks; VBNs, volume-based networks.

### Hamodynamic correlates of morphological WM networks

To examine the hamodynamic correlates of morphological WM networks, we used the communication model, a multilinear regression model based on simple dynamical processes (34), to predict the group-level mean hamodynamic coherence profile, with group-level mean similarity profile and communicational organization (shortest path length and communicability) derived from the morphological WM networks as predictors (35). We constructed the model at whole brain level and for each node. The responding variable and predictors were derived from the R-BOLD-fMRI and structural MRI data of 444 unrelated healthy participants in HCP dataset, respectively.

We observed the significant predictions of group-level mean hamodynamic coherence profiles with similarity profiles and communicational organization of the morphological WM networks at whole brain level (VBN: adjusted R-squared, *AR^2^* = 0.034, *P* = 0.004; DBN: *AR^2^* = 0.022, *P* = 0.017). Fig 5A top shows the spatial distribution of *AR^2^* values derived from node-wise multilinear predictions (VBN: - 0.061 - 0.192; DBN: −0.055 −0.417). For the VBN, no significant predictions were observed. For the DBN, significant predictions were for the left cerebral peduncle (*AR^2^* = 0.417, *P* < 0.001) and right posterior limb of internal capsule (*AR^2^* = 0.362, *P* < 0.001).

**Fig 5.**
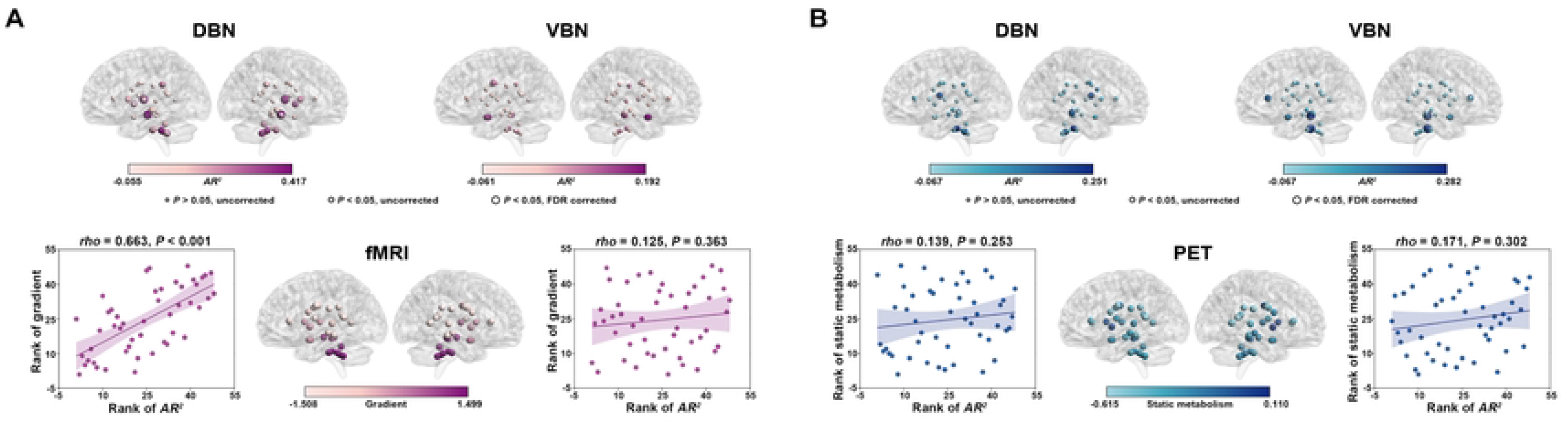
Hamodynamic and metabolic correlates of morphological WM networks. Top panels show spatial distribution of *AR^2^* values derived from node-wise multilinear predictions of group-level mean hamodynamic coherence (**A**) and metabolic synchronization (**B**) profiles with similarity profiles and communicational organization of the morphological WM networks. Bottom panels show spatial correlations of the *AR^2^* values with the first gradient derived from group-level mean hamodynamic coherence WM network (**A**) and static metabolic values (**B**). For the hamodynamic coherence, the DBN was able to predict the connectivity patterns of the left cerebral peduncle and right posterior limb of internal capsule, and the regional *AR^2^* values exhibited a significantly positive correlation with the gradient. For the metabolic synchronization, the VBN was able to predict the connectivity patterns of the pontine crossing tract and left cingulum hippocampus, and the regional *AR^2^* values were independent of static metabolic values. *AR^2^*, adjusted R-squared; DBN, deformation-based network; VBN, volume-based network; FDR, false discovery rate.

We further examined whether the nodal *AR^2^* values followed functional hierarchy of the WM networks. The first gradient accounted for 30.5% variance in the distribution of hamodynamic coherence across the WM regions. The gradient described an axis with gradual reduction from inferior to superior WM regions (Fig 5A bottom). We found that the nodal *AR^2^* values exhibited a significantly positive correlation with the gradient for the DBN (*rho* = 0.663, *P* < 0.001) but not VBN (*P* > 0.05) (Fig 5A bottom).

### Metabolic correlates of morphological WM networks

To examine the metabolic correlates of morphological WM networks, we used the communication model similar to the hamodynamic correlates. The responding variable and predictors were derived from the R-FDG-PET and structural MRI data of 26 healthy participants in MU dataset, respectively.

We observed the significant predictions of group-level mean metabolic synchronization profiles with similarity profiles and communicational organization of the VBNs but not the DBNs at whole brain level (VBN: *AR^2^* = 0.012, *P* = 0.017; DBN: *AR^2^* = 0.005, *P* = 0.411). Fig 5B top shows the spatial distribution of *AR^2^* values derived from node-wise multilinear predictions (VBN: −0.067 −0.282; DBN: −0.067 −0.251). For the VBN, significant predictions were found for the pontine crossing tract (*AR^2^* = 0.260, *P* < 0.001) and left cingulum hippocampus (*AR^2^* = 0.282, *P* < 0.001). No significant predictions were observed for the DBN.

We further linked the nodal *AR^2^* values with regional static metabolic values, and found no significant correlations for neither the DBN nor VBN (*P* > 0.05) (Fig 5B bottom).

### Chemoarchitectonic correlates of morphological WM networks

Mounting evidence indicates the existence of multiple neurotransmitters in WM tracts, such as serotonergic, glutamatergic, dopaminergic, GABAergic, purinergic, adrenergic and cholinergic signaling (36). Thus, we constructed a chemoarchitectonic WM network and used the communication model similar to the hamodynamic correlates to examine chemoarchitectonic correlates of the morphological WM networks. The responding variable and predictors were derived from the neurotransmitter receptor and transporter distributions in JuSpace dataset and structural MRI data of 444 unrelated healthy participants in HCP dataset, respectively.

We observed the significant predictions of group-level mean chemoarchitectonic covariance profiles with similarity profiles and communicational organization of the DBN but not the VBN at whole brain level (VBN: *AR^2^* = 0.022, *P* = 0.145; DBN: *AR^2^* = 0.056, *P* = 0.007). Fig 6 left shows the spatial distribution of *AR^2^* values derived from node-wise multilinear predictions (VBN: −0.067 −0.321; DBN: −0.067 −0.503). For the VBN, no significant predictions were observed. For the DBN, significant predictions were observed for the genu of corpus callosum (*AR^2^* = 0.296, *P* < 0.001), left cerebral peduncle (*AR^2^* = 0.318, *P* < 0.001), left anterior limb of internal capsule (*AR^2^* = 0.333, *P* < 0.001), left posterior limb of internal capsule (*AR^2^* = 0.227, *P* = 0.003), right corticospinal tract (*AR^2^* = 0.344, *P* < 0.001), right inferior cerebellar peduncle (*AR^2^* = 0.503, *P* < 0.001) and right anterior limb of internal capsule (*AR^2^* = 0.322, *P* < 0.001).

**Fig 6.**
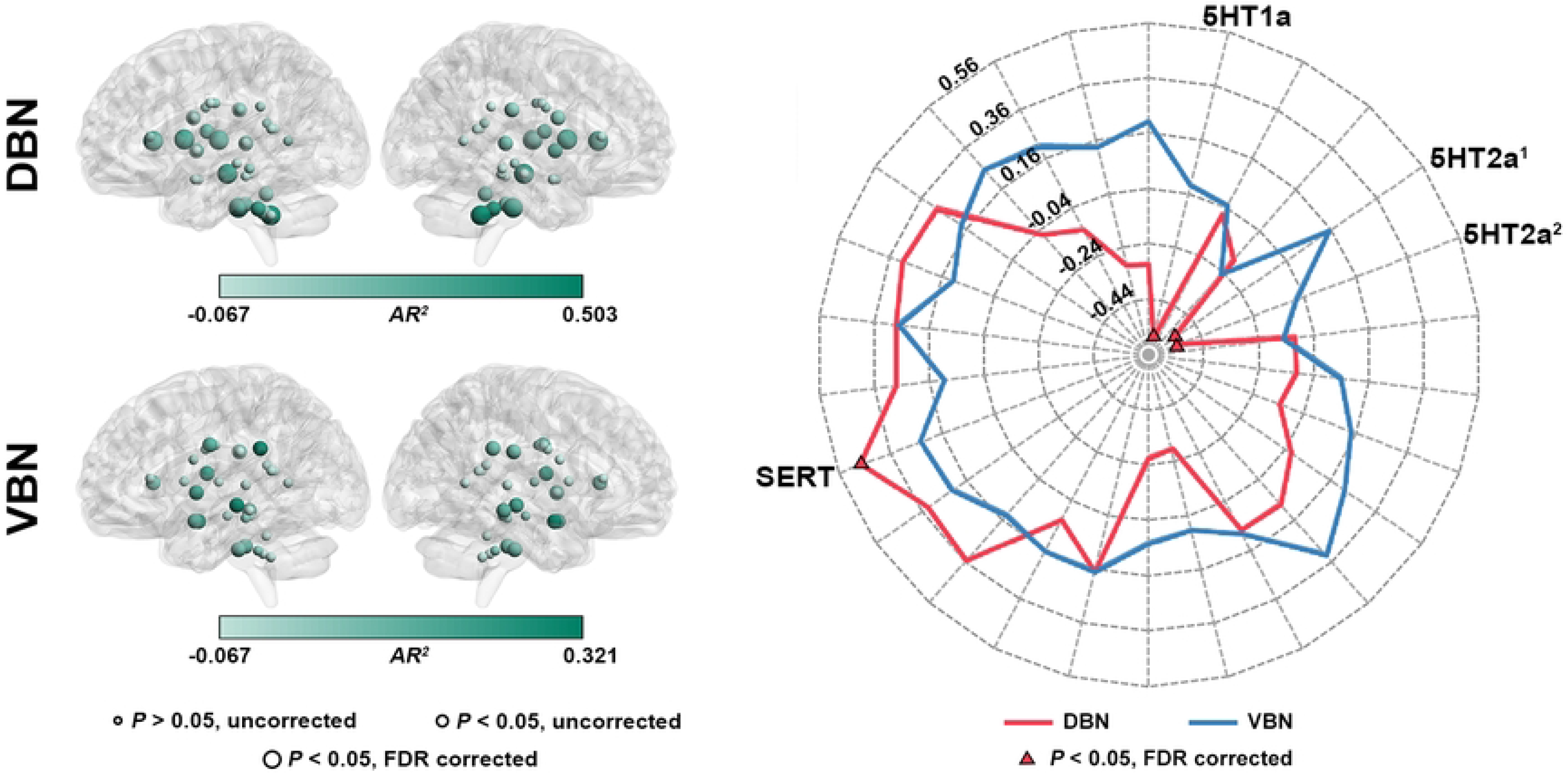
Chemoarchitectonic correlates of morphological WM networks. For the DBN, similarity profiles and communicational organization of the morphological WM networks were able to predict the patterns of chemoarchitectonic covariance for the genu of corpus callosum, left cerebral peduncle, left anterior limb of internal capsule, left posterior limb of internal capsule, right corticospinal tract, right inferior cerebellar peduncle and right anterior limb of internal capsule. Moreover, the regional *AR^2^* values were correlated with regional neurotransmitter density of SERT with ^11^C-MADAM tracer, 5HT1a with ^11^C-CUMI tracer and 5HT2a with ^18^F-ALTANSERIN tracer and ^11^C-CIMBI tracer. No significant predictions were observed for the VBN. *AR^2^*, adjusted R-squared; DBN, deformation-based network; VBN, volume-based network; FDR, false discovery rate. ^1^The 5HT2a maps were collected using ^18^F-ALTANSERIN tracer. ^2^The 5HT2a maps were collected using ^11^C-CIMBI tracer.

We further examined the associations of the nodal *AR^2^* values with regional neurotransmitter density. For the VBN, no significant correlations were observed (*P* > 0.05, FDR corrected). For the DBN, the nodal *AR^2^* values exhibited a significantly positive correlation with the density of SERT with ^11^C-MADAM tracer (*rho* = 0.476, *P* = 0.002), and negative correlations with the density of 5HT1a with ^11^C-CUMI tracer (*rho* = −0.574, *P* = 0.002) and 5HT2a with ^18^F-ALTANSERIN tracer (*rho* = −0.529, *P* = 0.002) and ^11^C-CIMBI tracer (*rho* = −0.533, *P* = 0.002) (Fig 6 right).

### Genetic correlates of morphological WM networks

To examine the genetic correlates of morphological WM networks, we used the communication model similar to the hamodynamic correlates. The responding variable and predictors were derived from the brain-wide transcriptional profiles and structural MRI data in AHBA dataset, respectively.

We observed the significant predictions of group-level mean gene co-expression profiles with similarity profiles and communicational organization of the VBN but not the DBN at whole brain level were observed (VBN: *AR^2^* = 0.045 *P* = 0.061; DBN: *AR^2^* = 0.009, *P* = 0.439). Fig 7A top shows the spatial distribution of *AR^2^* values derived from node-wise multilinear predictions (VBN: −0.080 −0.288; DBN: −0.079 −0.281). For the VBN, no significant predictions were observed. For the DBN, a significant prediction was observed for the right uncinate fasciculus (*AR^2^* = 0.281, *P* < 0.001).

**Fig 7.**
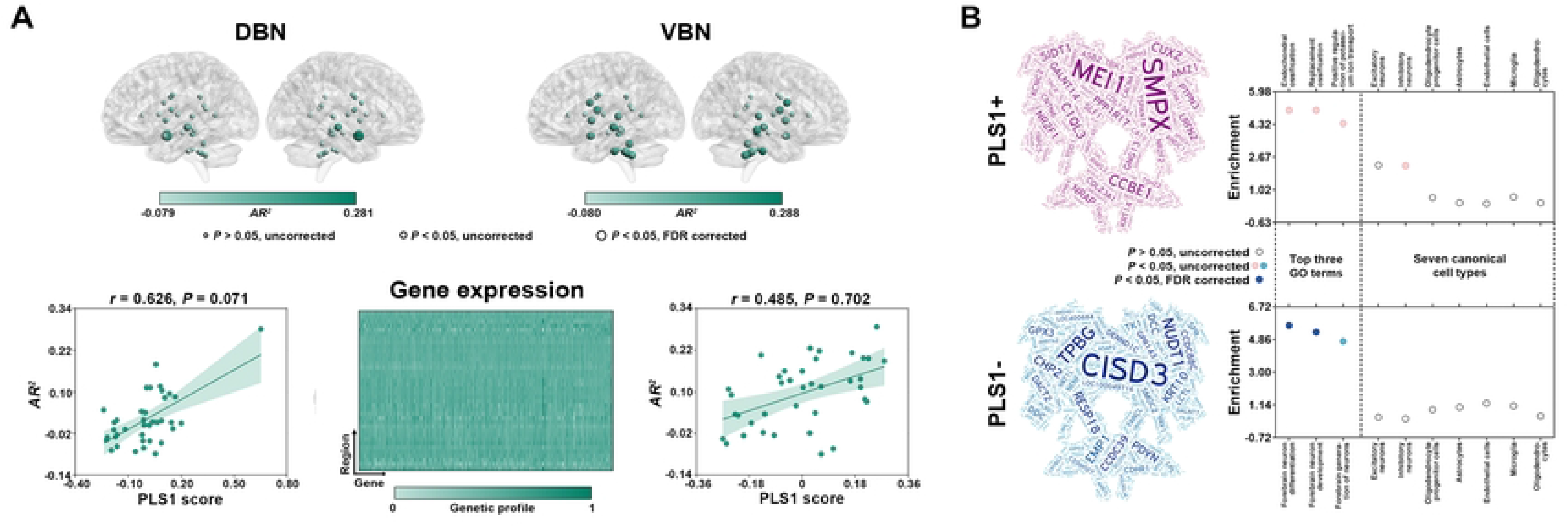
Genetic correlates of morphological WM networks. **A**) The spatial distribution of regional *AR^2^* values derived from node-wise multilinear predictions of gene co-expression with group-level mean similarity profiles and communicational organization of the morphological WM network. The DBN was able to predict the patterns of gene co-expression for the right uncinate fasciculus. Moreover, the distribution of regional *AR^2^* values can be explained by gene spatial expression. **B**) Functional annotations of genes determining the distribution of regional *AR^2^* values. The PLS1-genes were enriched in two biological processes of “forebrain neuron differentiation” and “forebrain neuron development” but did not exhibit susceptibility to certain cell types. No results were found for the PLS1+ genes. *AR^2^*, adjusted R-squared; DBN, deformation-based network; VBN, volume-based network; FDR, false discovery rate; GO, gene ontology; PLS, partial least-squares.

We further linked the nodal *AR^2^* values with regional gene expression. No significant correlations were observed for the VBN (*r* = 0.485, *P* = 0.702). For the DBN, a significantly positive correlation was observed (*r* = 0.626, *P* = 0.071) with 1442 genes strongly contributing to this correlation (the first positive component of the PLS, PLS1+: 837; the first negative component of the PLS, PLS1-: 605) (Fig 7A bottom). Gene ontology (GO) enrichment analysis revealed that the PLS1-genes were enriched in two biological processes of “forebrain neuron differentiation” and “forebrain neuron development”, while no results were observed for the PLS1+ genes. Neither the PLS1+ nor PLS-genes exhibited susceptibility to certain cell types (*P* > 0.05) (Fig 7B).

### Clinical application of morphological WM networks

Based on the MS and NMOSD multicentric dataset with 208 MS patients, 200 NMOSD patients, and 228 healthy controls (HCs), the clinical correlates of morphological WM networks were explored by examining disease-related alterations in interregional morphological similarity, associations of altered morphological similarity with clinical variables, and classification and differentiation of MS and NMOSD.

#### Altered morphological WM networks in MS and NMOSD

A total of 106 and 10 edges were identified to exhibit significant group main effects on interregional morphological similarities in the VBNs and DBNs, respectively (*P* < 0.05, FWE corrected). Post hoc comparisons revealed that the edges exhibited MS-specific alterations (VBNs: 99; DBNs: 8), common alterations to MS and NMOSD (VBNs: 5; DBNs: 2) or opposite alterations between MS and NMOSD (VBNs: 2) (Table S1; Fig 8A).

**Fig 8.**
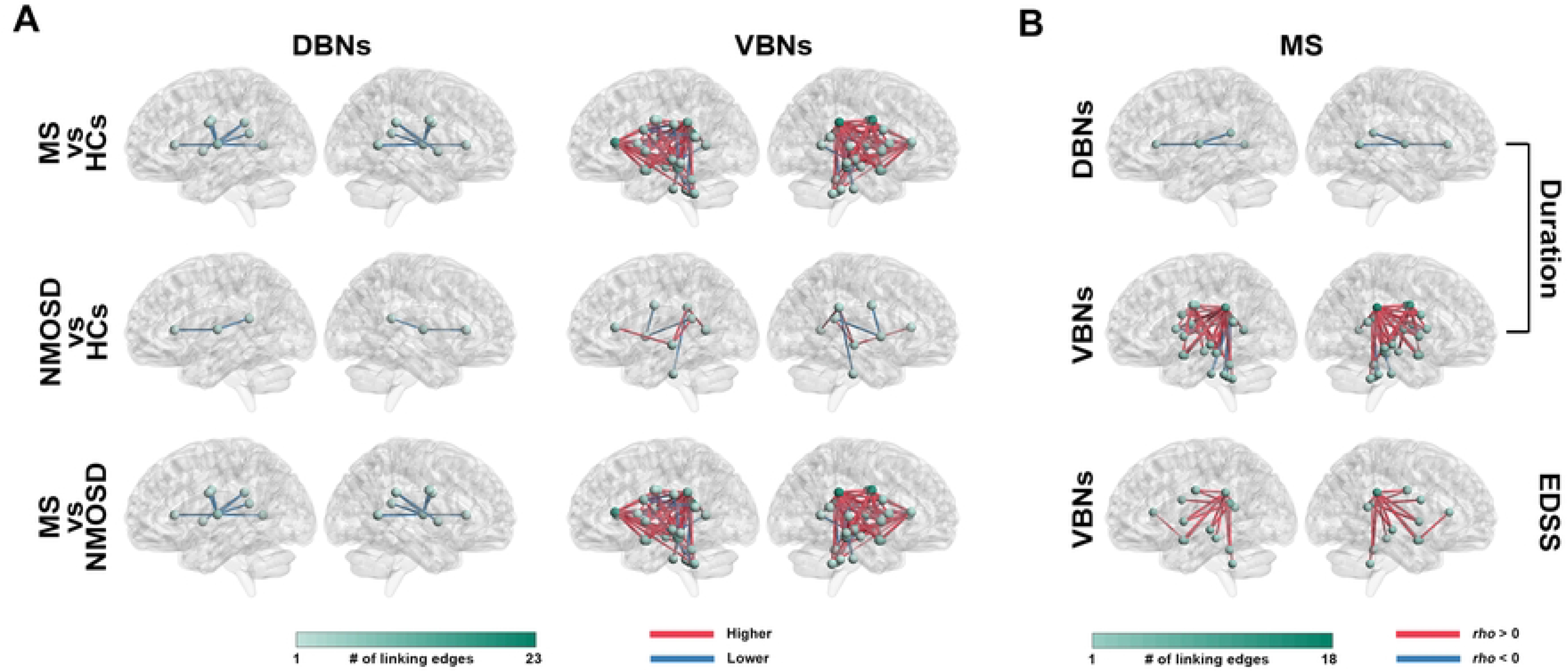
Disrupted morphological WM networks in MS and NMOSD. **A**) Connections showing significant between-group differences in morphological similarities. Both diseases were associated with disrupted morphological WM networks, in particular for MS in the VBNs. **B**) Clinical correlations of disrupted connections in MS. Out of the connections showing altered morphological similarities in MS, many showed significantly correlations with disease duration and Expanded Disability Status Scale scores of the patients. Notably, the connections were mainly linked to the splenium of corpus callosum and corona radiate. DBNs, deformation-based networks; VBNs, volume-based networks; EDSS, Expanded Disability Status Scale; FDR, false discovery rate; HCs, healthy controls; MS, multiple sclerosis; NMOSD, neuromyelitis optica spectrum disorders.

#### Associations of altered morphological WM networks with clinical variables

Among the 106 edges in the VBNs showing altered morphological similarities in MS, 50/5 edges showed significantly positive/negative correlations with disease duration, and 21 edges showed significantly positive correlations with Expanded Disability Status Scale scores of the patients (*P* < 0.05, FDR corrected). Out of the 10 edges in the DBNs showing altered morphological similarities in MS, 4 edges exhibited significantly negative correlations with disease duration of the patients (*P* < 0.05, FDR corrected) (S1 Table; Fig 8B). For the edges in the VBNs and DBNs showing altered morphological similarities in NMOSD, no significant correlations were found with any clinical variables of the patients (*P* > 0.05, FDR corrected).

#### Classification and differentiation of MS and NMOSD

The VBNs distinguished the three groups from each other (MS vs HCs: mean accuracy = 77.3%, *P* < 0.001; NMOSD vs HCs: mean accuracy = 62.2%, *P* < 0.001; MS vs NMOSD: mean accuracy = 74.5%, *P* < 0.001), while the DBNs failed to distinguish NMOSD from HCs (MS vs HCs: mean accuracy = 69.6%, *P* < 0.001; MNOSD vs HCs: mean accuracy = 54.4%, *P* = 0.104; MS vs NMOSD: mean accuracy = 60.5%, *P* < 0.001).

## Discussion

In this study, we constructed volume- and deformation-based single-subject morphological WM networks using structural MRI data, and systematically investigated their topological organization, TRT-reliability, phenotypic association, heritability, functional relevance, neurobiological substrate and clinical value by combining multimodal and multiscale data. We found that the morphological WM networks exhibited nontrivial topological organization, presented good to excellent TRT reliability, accounted for phenotypic interindividual differences, were under genetic control, shaped WM hamodynamic coherence and metabolic synchronization, were determined by gene co-expression due to genes enriched in the forebrain neuron development and differentiation, predicted chemoarchitectonic covariance by means of serotonergic system-related receptors and transporters, and were able to help diagnose and differentiate MS and NMOSD. Overall, these findings deepen our understanding of the roles and origins of morphological WM networks, and provide convincing evidence for the usage of morphological WM networks for individualized research in health and disease.

### Specifically organized morphological WM networks

Numerous studies have demonstrated the nontrivial organizational principles in multimodal gray matter networks, such as small-worldness, modularity and rich-club organization (3, 5, 37, 38). In recent years, the nontrivial organizational principles are also found to exist in functional WM networks (15). Here, our findings further indicated that morphological WM networks were specifically organized in an optimized manner to support efficient information spreading and processing, which may be structural substrates of the wiring patterns in functional WM networks. Moreover, the optimized configurations were consistently observed regardless of the choice of morphological index and whether spatial smoothing was performed. These findings suggest that the optimized configurations are intrinsic characteristics of morphological WM networks, presumably as a consequence of evolution by natural selection. Nevertheless, it should be noted that the DBNs and VBNs exhibited low correlations in their wiring patterns, suggesting that they are complementary to each other in mapping morphological WM networks. The low correlations may be due to fundamental differences in the nature between volume (reflecting a composite of thickness, area and folding) and deformation (reflecting local tissue expansion or contraction), and/or different neurobiological substrates between the VBNs and DBNs (see our discussion below).

### TRT reliable morphological WM networks

High TRT reliability is a prerequisite for a new method or measure before its use as potential clinical biomarkers. Here, we found that most interregional morphological WM similarities exhibited good to excellent TRT reliabilities regardless of the scanning intervals. These findings indicate that morphological WM networks can serve as a promising approach to reliably examine effects of interest on WM architecture. It should be noted that the TRT reliabilities of the morphological WM networks were obviously higher than those reported for functional WM networks (12). Therefore, relative to functional WM networks, morphological WM networks are a preferable choice for study wiring patterns of WM from the perspective of TRT reliability. Furthermore, we found that the TRT reliabilities of morphological WM networks depended on the choice of morphological index used for network construction (VBNs > DBNs) and can be further improved by performing spatial smoothing. These findings provide guidance on determining reliable analytical strategies for morphological WM networks and are consistent with our previous findings from morphological gray matter networks (25, 39). Overall, we propose morphological WM networks as a reliable approach, which can be used to examine the association between their variability and phenotypic interindividual differences.

### Behavioral and cognitive relevance of morphological WM networks

Numerous previous studies have demonstrated tight associations of multimodal gray matter networks with behavior and cognition (3, 40–43). For functional WM networks, a preliminary study found that the small-world organization was correlated with fluid intelligence in healthy participants (15). Here, by combining multiple behavioral and cognitive domains with multivariate approaches, we showed that morphological WM networks were able to explain interindividual variance and predict individual performance in specific behavioral and cognitive domains. These findings are consistent with our previous study of morphological gray matter networks (41). Specifically, both the VBNs and DBNs captured interindividual differences in the Motor domain. These findings sound plausible given the crucial roles of WM tracts in conveying motor information (44) and previous findings that WM volume and deformation were related to motor performance (45, 46). In addition to the Motor domain, the VBNs were related to individual performance in the Cognition domain. This is consistent with previous studies showing the relationship between WM volume and cognition (47–49). Therefore, our findings suggest that relative to the DBNs, the VBNs can be preferentially used to study neural correlates of cognition and brain diseases accompanied by cognitive dysfunction. Notably, the behavioral and cognitive associations of morphological WM networks were examined with all connections as a whole. Given previous findings from gray matter networks that different sets of highly interconnected nodes (i.e., modules) play domain-specific roles in cognition (1, 50, 51), it is thus an interesting topic to explore behavioral and cognitive relevance of morphological WM networks in the context of modular architecture.

### Functional correlates of morphological WM networks

How structural connectivity supports and shapes functional connectivity in the brain is a fundamental question in systems neuroscience. In this study, we found that morphological WM networks were predictive of regional connectivity profiles derived from temporal fluctuations in hemodynamic activity and glucose metabolism, indicating structural-functional coupling of WM networks. Interestingly, the DBNs accounted for regional patterns of hamodynamic coherence, while the VBNs largely determined regional profiles of metabolic synchronization. The dissociation of structural-functional coupling further highlights potentially distinct origins between the DBNs and VBNs. Moreover, we found that the extents to which the DBNs explained variance of regional profiles of hamodynamic coherence followed functional hierarchy of WM networks. This is consistent with previous findings from gray matter networks that the relationship between fiber tractography and hamodynamic coherence followed functional network hierarchy (35). It thus seems that the constraint on structural-functional coupling from functional hierarchy is a fundamental rule of human brain regardless of gray matter and WM networks. It is worth noting that in contrast to the well characterized functional hierarchy of gray matter networks (52–55), significant gaps remain in our understanding of the functional hierarchy derived from WM networks with respect to its exact spatial pattern, developmental trajectory and associations with other data modalities (e.g., gene expression and chemoarchitecture). Elucidating these issues can deepen our understanding of functional organization of WM networks, thereby helping explain the structural-functional coupling observed in this study. Overall, our findings provide strong evidence supporting that morphological WM networks determine, to some extent, the wiring patterns of functional WM networks.

### Neurobiological substrates of morphological WM networks

In this study, we examined neurobiological substrates of morphological WM networks by linking them with genetic expression and chemoarchitecture. As for genetic bases, we first calculated edgewise heritability to show the extent to which the morphological WM networks were heritable. We found that both the VBNs and DBNs were under genetic control with varying degrees across edges. Moreover, the heritability was significantly higher for the VBNs than DBNs. These findings indicate the important roles of genes in determining the morphological WM networks, particularly for the VBNs. Intriguingly, we found significantly positive correlations for the VBNs between the edge’s heritability and contribution to accounting for interindividual differences in the Motor and Cognition domains. These correlations indicate that highly heritable edges in the VBNs are strongly correlated with individual motor and cognitive abilities, and thus the VBNs may serve as potential intermediate phenotype to link gene and behavior and cognition. In addition to the heritability analysis, we examined the relationships between morphological and transcriptional WM networks. We found that the DBNs predicted gene co-expression with the spatial pattern determined by a specific set of genes enriched in the forebrain neuron development and differentiation. The forebrain neuron development and differentiation support the progression of a neuron in the forebrain from its initial commitment to its fate, and finally to the fully functional differentiated cell. A previous study found that initial axon extension starts adjacent to a focus of microtubule polymerization within the cell body and the microtubules grow into the developing axon (56). Thus, the structural organization of the cell may establishes the initial site and direction of axonal extension at the time of neuronal differentiation (57). Based on these findings, our results suggest that the forebrain neuronal progression plays an important role in the formation of the DBNs by modulating the growth of WM. It should be noted that despite under genetic control, the VBNs failed to predict gene co-expression, and no genes were identified to account for the spatial pattern of predictive performance. This may be attributed to that the VBNs are relatively independent of the genes used in the current study.

In addition to the genetic basis, we examined the relationships between morphological and chemoarchitectonic WM networks. We found that the DBNs could predict chemoarchitectonic covariance between WM regions, suggesting the chemoarchitectonic basis of morphological WM networks. The chemoarchitectonic basis was also found for morphological gray matter networks in our previous study (Li, 2022). Thus, it seems that chemoarchitecture may be a common source of morphological brain networks for both gray matter and WM. Specifically, we found that the morphological-chemoarchitectonic relationship across WM regions was correlated with the distributions of several serotonergic system-related receptors (5HT1a and 5HT2a) and transporter (SERT). Serotonin has a diverse range of physiological roles, such as cell growth and differentiation (58), neuronal development (59) and neuronal signaling pathways. For WM, previous studies showed that the activation of 5HT1a and 5HT2a modulated the axonal excitability in rat spinal dorsal columns (60, 61). In humans, the relationship between serotonin and WM was also increasingly reported. For instance, infants exposed to selective serotonin reuptake inhibitor were found to show increased WM structural connectivity (62). It should be noted that the physiological functions of neurotransmitter signaling in WM are debatable, and future work is needed for deeper understanding the exact roles of chemoarchitecture in shaping morphological WM networks. Overall, our findings suggest that morphological WM networks are a neurobiologically meaningful approach to provide an avenue for linking WM organization with molecular biological processes.

### Altered morphological WM networks in MS and NMOSD

To test clinical value of our method, we applied the morphological WM networks to MS and NMOSD, both of which are characterized by multifocal areas of WM lesions. We found that both diseases exhibited abnormalities (increases and decreases) in interregional WM similarities, particularly in the VBNs. These findings suggest that the MS and NMOSD can be seen as disorders of morphological WM dysconnectivity. Nevertheless, much more alterations were observed in MS than NMOSD, indicative of more serious morphological WM dysconnectivity in MS. Moreover, many of the alterations in MS were correlated with disease duration and Expanded Disability Status Scale scores of the patients, highlighting the potential of the morphological WM dysconnectivity in monitoring disease progression of MS. Interestingly, both the alterations and correlations in MS were mainly involved in edges linking the splenium of corpus callosum and corona radiate, where the earliest WM atrophy occurred in MS (63). These findings imply the important roles of the splenium of corpus callosum and corona radiate in understanding the pathophysiology of MS. In the future, it is interesting to explore the relationship between WM atrophy and dysconnectivity as MS progresses. For NMOSD, fewer alterations were observed, and most of the alterations were also identified in MS in the same direction. These common alterations provide novel insights into shared neural mechanisms between MS and NMOSD from the perspective of morphological WM dysconnectivity. Interestingly, two edges in the VBNs exhibited opposite patterns in the alterations between MS and NMOSD. These results together with the findings that numerous edges in the VBNs were altered in MS but intact in NMOSD collectively suggest that relative to the DBNs the VBNs may be a better choice in uncovering diagnosis-specific biomarkers in MS and NMOSD. This speculation was supported by our classification analyses revealing higher accuracies for the VBNs than DBNs in distinguishing the three groups from each other. Overall, our preliminary attempts demonstrate the clinical value of morphological WM networks in helping diagnose and differentiate MS and NMOSD. Notably, the classification accuracies were not that high, presumably due to the simple classification models used in this study, which can be further improved by using more sophisticated machine learning techniques and optimizing model parameters in the future (64, 65).

### Limitation and future directions

First, morphological WM networks were constructed separately with volume and deformation maps, both of which were widely used to characterize WM architecture. In addition to these two measures, whether our method could be extended to other WM features (e.g., fractional anisotropy) is an open question, and studies on this topic may help more comprehensively understand structural wiring patterns between WM regions. Second, effects of different analytical choices, including brain parcellation, connectivity estimation, and thresholding method have been well documented in the literature for multimodal gray matter networks (25, 66–71). Intuitively, these factors will also affect topological characterization of morphological WM networks, which should be extensively studied in future work. Third, the chemoarchitectonic WM network used for examining chemoarchitectonic correlates of morphological WM networks was constructed with publicly available data from various studies. This may result in an underestimation of the relationship between chemoarchitectonic and morphological WM networks. Future work is needed to validate our results by collecting related data from the same cohort of participants. Finally, we only tested whether morphological WM networks were predictive of other modalities of WM networks. A recent study found that different predictors derived from diffusion MRI-based structural networks predicted hamodynamic coherence profiles of specific regions (72). Thus, a more detailed mapping of the relationship between morphological and other modalities of WM networks can be obtained by identifying region-wise optimal predictors and disclosing how the optimal predictors vary across regions.

### Materials and Methods Participants and data acquisition

#### HCP dataset

Of 1113 participants with T1-weighted structural images in the HCP dataset, a total of 444 unrelated healthy participants (age, 22 - 35 years; 205 males and 239 females) and 217 pairs of MZ and DZ twins (age, 22 - 35 years; 174 males and 260 females) were included in this study. All MRI images were acquired using a customized 3T scanner at Washington University in St. Louis. The structural images were acquired using a magnetization-prepared rapid gradient-echo sequence with the following parameters: repetition time (TR) = 2,400 ms, echo time (TE) = 2.14 ms, flip angle (FA) = 8°, 256 slices, matrix = 320 × 320, field of view (FOV) = 224 × 224 mm^2^, slice thickness/gap = 0.7/0 mm, and voxel size = 0.7 × 0.7 × 0.7 mm^3^. The R-BOLD-fMRI data were obtained using a multiband echo-planar imaging sequence: TR = 720 ms, TE = 33 ms, FA = 52°, 72 slices, matrix = 104 × 90, FOV = 208 × 180 mm^2^, slice thickness/gap = 2/0 mm, and voxel size = 2 × 2 × 2 mm^3^. The R-BOLD-fMRI data were acquired in two sessions for each participant on consecutive days and each session consisted of two runs with left-to-right and right-to-left phase encoding protocols. The length of each R-BOLD-fMRI scan was 14.4 min (1,200 volumes). In this study, only the R-BOLD-fMRI data with the left-to-right phase encoding in the first session were used.

In addition to the imaging data, the HCP dataset included a broad range of behavioral and cognitive measures that were evaluated mainly via the NIH Toolbox Assessment of Neurological and Behavioral function. For the initial 581 items of behavioral and cognitive measures, we performed a multistep screening strategy in terms of our research purpose. First, items related to health and family history, psychiatric and life function and substance use were excluded. Items resulting from magnetoencephalogram and functional MRI tasks (e.g., accuracy and reaction time) were also deleted. For the remaining items, we further ruled out those that had the same values or were missing for more than 80% of the participants since they may fail to sufficiently capture interindividual differences. Furthermore, for items with unadjusted and adjusted values by age, original scores were used since using age-adjusted versions has little effect on the relationships between brain networks and behavior (73). In addition, if total scores were provided for several items, the total scores were retained instead of individual items. After these procedures, a total of 60 items remained in this study, which were categorized into 6 behavioral and cognitive domains including Alertness, Cognition, Emotion, Motor, Personality, and Sensory. Based on the 60 items, participants who did not complete any tests in at least one behavioral and cognitive domain were excluded from this study. To further exclude the effects of outliers, values that were 3.29 standard deviations above or below the mean for each item were replaced by the mean ± 3.29 standard deviations. Finally, the items were averaged within each domain after standardizing each item to make the scale comparable (Z-transformation across participants).

#### HNU dataset

The HNU dataset included 30 participants (age, 20 - 30 years; 15 males and 15 females) that were scanned ten times over one month with one scan every three days. None of participants had a history of neurological or psychiatric disorders, substance abuse or head injury with loss of consciousness. T1-weighted structural images were obtained on a GE MR750 3.0 Tesla scanner (GE Medical Systems, Waukesha, WI) using a Fast Spoiled Gradient echo sequence: TR = 8.1 ms, TE = 3.1 ms, inversion time (TI) = 450 ms, FA = 8°, FOV = 256 × 256 mm^2^, matrix = 256 × 256, and voxel size = 1 × 1 × 1 mm^3^, slices = 176.

#### SWU dataset

The SWU dataset contained 580 participants at baseline scan (age, 17 - 27 years; 60 males and 60 females), some of whom completed the second (240) and third (228) scans. The average time intervals were 304.14, 515, and 817.87 days between the first and second scans, second and third scans, and first and third scans, respectively. All participants had no history of neurological or psychiatric disorders. T1-weighted structural images were obtained on a 3T Siemens Trio MRI scanner using the magnetization-prepared rapid gradient-echo sequence: TR = 1,900 ms, TE = 2.52 ms, TI = 900 ms, FA = 9°, FOV = 256 × 256 mm^2^, matrix = 256 × 256, slice thickness = 1.0 mm, slices = 176, and voxel size = 1 × 1 × 1 mm^3^. In this study, a total of 121 participants who completed all three scans were included.

#### MU dataset

The MU dataset contained 27 participants who underwent a 95-minute simultaneous MR-PET scan in a Siemens (Erlangen) 3 Tesla Biograph molecular MR scanner (Syngo VB20 P). One participant was excluded from further analyses due to the poor performance of PET-MRI co-registration (age, 18 - 23 years; 7 males and 19 females). All participants had no history of diagnosed Axis-1 mental illness, diabetes, or cardiovascular illness. Over the course of the scan, [18-F] fluorodeoxyglucose (average dose 233MBq) was infused at a rate of 36 mL/hr using a BodyGuard 323 MR-compatible infusion pump (Caesarea Medical Electronics, Caesarea, Israel). The detailed protocols of R-FDG-PET collection were listed elsewhere (29). T1-weighted structural images were acquired on a 3T Siemens Biograph molecular MRI scanner using the magnetization-prepared rapid gradient-echo sequence: TR = 1,640 ms, TE = 2.34 ms, slices = 176, FOV = 256 × 256 mm^2^, FA = 8°, and voxel size = 1 × 1 × 1 mm^3^.

#### JuSpace dataset

The JuSpace dataset provided 27 neurotransmitter receptor and transporter maps from different studies on healthy participants including 5HT1a, 5HT1b, 5HT2a, 5HT4, CB1, D1, D2, DAT, ^18^F-DOPA, GABAa, mGluR5, MU, NAT, SERT and VAChT. The D2 map derived from the raclopride tracer was excluded from further analysis because this tracer had unreliable binding in the cortex. Thus, a total of 26 neurotransmitter receptor and transporter maps were used in this study.

#### AHBA dataset

The AHBA dataset is a publicly available online resource of brain-wide transcriptomic information and multimodal MRI obtained from six healthy adult human donors (age, 24 - 57 years; 5 males and 1 female) with no known history of neuropathological or neuropsychiatric disease (31). Specifically, transcriptional activity was recorded for 20,737 genes from 3,702 spatially distinct tissue samples that covered almost the entire brain. The tissue samples were collected from the left hemisphere for 4 donors and both hemispheres for 2 donors. Prior to dissection, T1-weighted structural MRI images were acquired for each donor on a 3T Siemens Magnetom Trio scanners (Erlangen, Germany) using the MPRAGE sequence: TR = 1,900 ms, TI = 900 ms, and FA = 9°. Other imaging parameters differed among the donors. For more details, see http://human.brain-map.org/.

#### MS and NMOSD multicentric dataset

The MS and NMOSD multicentric dataset contained 208 MS patients, 200 NMOSD patients, and 228 HCs (age, 18 - 65 years; 104 males and 124 females in HCs, 73 males and 135 females in MS, 25 males and 175 females in NMOSD). Details on the inclusion and exclusion criteria of patient selection and imaging parameters were provided elsewhere (32). This study was approved by the institutional review board of corresponding hospitals, and written informed consent was obtained from each participant.

### Preprocessing of structural images

All structural images underwent the same standard preprocessing pipeline using the CAT12 (http://www.neuro.uni-jena.de/cat/) based on the SPM12 package (https://www.fil.ion.ucl.ac.uk/spm/software/spm12). Briefly, each structural image was first segmented into gray matter, WM and cerebrospinal fluid using an adaptive Maximum A Posterior technique. The resulting WM maps were then normalized to the standard Montreal Neurological Institute (MNI) space using a geodesic shooting approach (74). WM volume was calculated by modulating the WM maps using Jacobian determinants derived from the spatial normalization. Finally, a WM volume map and deformation map (i.e., Jacobian determinants) were obtained for each structural image. Of note, the Segment Longitudinal Data module in the CAT12 was used for tissue segmentation of structural images in the SWU and HNU datasets.

### Spatial smoothing

Spatial smoothing is frequently used to increase the signal-to-noise ratio, and improve interindividual anatomical correspondence for voxel-based morphology analysis. However, it may introduce spurious local connectivity between spatially adjacent regions. In our previous study, we found that spatial smoothing significantly affected the wiring patterns and topological organization of morphological brain networks between gray matter regions, and the implementation of spatial smoothing increased the TRT reliability (39). To evaluate the effects of spatial smoothing on morphological WM networks, individual WM volume and deformation maps with (Gaussian kernel with 8-mm full width at half maximum) and without spatial smoothing were separately used to construct morphological WM networks.

### Construction of morphological WM networks

Morphological WM networks were constructed using our previous reliable approaches for depicting single-subject morphological brain networks between gray matter regions (25, 39). First, a WM atlas from Johns Hopkins University (24) was used to divide brain WM into 48 ROIs. Then, values of WM volume and deformation were separately extracted from all voxels within each ROI, and used to derive regional probability distribution functions (PDFs). Afterwards, interregional morphological similarity was estimated as:

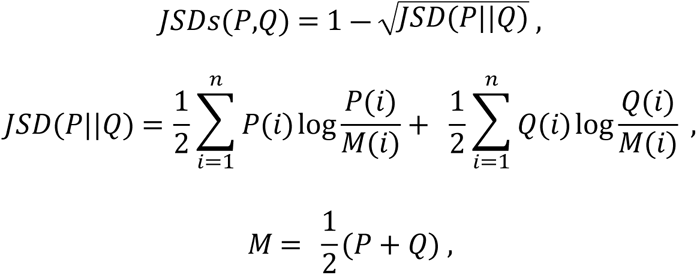

where *P* and *Q* denote regional PDFs, and *n* denotes the number of sample points (2^8^in this study) (39). The value of JSDs indicates the extent to which two PDFs are similar (0 indicating completely different and 1 exactly the same). Finally, for each structural image a morphological WM network was obtained under each combination of morphological index (volume vs deformation) and spatial smoothing (yes vs no).

### Effects of analytical strategies on morphological WM networks (HCP dataset, unrelated participants)

To explore the effects of morphological index and spatial smoothing on the morphological WM networks, we first calculated the Spearman correlation between the group-level mean VBN and DBN and the Spearman correlation of the group-level mean morphological WM networks between data with and without spatial smoothing. Significance levels of the correlations were estimated through Moran spectral randomization to account for spatial autocorrelation (10,000 times) (75). Furthermore, we performed two-way repeated ANOVA on the group-level mean morphological similarity of all edges in the morphological WM networks. For significant effects, post hoc comparisons were further performed with paired *T* tests.

### Topological analysis of morphological WM networks (HCP dataset, unrelated participants)

#### Threshold selection

Before topologically characterizing the morphological WM networks derived above, a sparsity-based thresholding procedure was first used to convert the morphological WM networks to binary networks. Sparsity is defined as the ratio of the number of actual edges divided by the maximum possible number of edges in a network. By applying subject-specific thresholds to the morphological WM networks, the sparsity-based thresholding procedure ensures the same number of edges across participants and visits under different analytical strategies. Given the lack of a definitive standard for choosing a single sparsity, a consecutive sparsity range of [0.09 0.3] with an interval of 0.02 was used in this study. The sparsity range was determined to guarantee that the resulting binary networks are sparse and estimable for the small-world attributes (66, 76, 77). To ensure that there were no isolated nodes or multiple connected components in the resulting binary networks, a minimum spanning tree algorithm (78) was further integrated into the sparsity-based thresholding procedure. For the resulting binary networks, several graph-based network measures were calculated with the GRETNA toolbox (79). Detailed formulas and interpretations of the network measures can be found elsewhere (80–82).

#### Small-worldness and rich-club organization

To test whether the morphological WM networks were non-randomly organized, we calculated the small-world attributes (clustering coefficient, *C_p_*, and characteristic path length, *L_p_*) and rich-club coefficient, *Φ* for each WM network at each sparsity. These global measures were further normalized by the corresponding mean of 100 matched random networks, which were generated with a topological rewiring algorithm to preserve the same degree distribution as the real networks (83). Typically, a small-world network should exhibit a normalized clustering coefficient > 1 and a normalized characteristic path length ∼ 1 (77), and a network with rich-club organization should exhibit a normalized rich-club coefficient > 1 (82).

#### Degree distribution and hubs

To characterize the roles of individual nodes in the morphological WM networks, we calculated nodal degree for each WM network at each sparsity. After averaging nodal degree across participants and over the entire sparsity range, different models (power law, exponential, and exponentially truncated power law) were used to fit the degree distribution of the morphological WM networks, and regions with a Z-transformed mean nodal degree > 1 were identified as hubs.

### TRT reliability of morphological WM networks (HNU dataset and SWU dataset)

The *ICC* (84) was used to quantify both short-term (HNU dataset) and long-term (SWU dataset) TRT reliability of morphological WM networks. Formally, for each edge in the morphological WM networks under all combinations of morphological index and spatial smoothing, the *ICC* was calculated as:

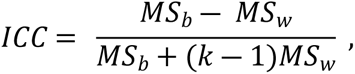

where *MS_b_* is the mean square of between-subject variance, *MS_w_* is the mean square of within-subject variance, and *k* denotes the number of repeated observations per participant (10 for the HNU dataset and 3 for the SWU dataset). *ICC* is close to 1 for reliable measures and 0 otherwise. In accordance with our previous studies (25, 39), the TRT reliability was categorized as poor (*ICC* < 0.25), low (0.25 < *ICC* < 0.4), fair (0.4 < *ICC* < 0.6), good (0.6 < *ICC* < 0.75) and excellent (0.75 < *ICC* < 1).

### Effects of analytical strategies on TRT reliability of morphological WM networks (HNU dataset and SWU dataset)

To examine the effects of morphological index and spatial smoothing on the TRT reliability of morphological WM networks, we performed two-way repeated ANOVA separately on the short-term and long-term reliabilities of all edges in the morphological WM networks. For significant effects, post hoc comparisons were further performed with paired *T* tests.

Given that the implementation of spatial smoothing significantly increased both short-term and long-term reliabilities of the morphological WM networks (see Results), only the VBNs and DBNs derived from spatially smoothed data were used for subsequent analyses.

### Behavioral and cognitive correlates of morphological WM networks (HCP dataset, unrelated participants)

Our previous study found that morphological brain networks of gray matter were able to explain interindividual variance and predict individual performance in specific behavioral and cognitive domains (41). Here, we further examined whether this ability was shared by morphological WM networks. Specifically, PLS regression was used to examine the relationship between the morphological WM networks and behavioral and cognitive data at each domain. In the PLS regression model, the response variable was behavioral and cognitive data at a certain domain, and the predictor variables were all interregional morphological similarities. The PLS1 was the linear combination of the interregional morphological similarities that exhibited the strongest correlation with the behavioral and cognitive data. Significance levels of the correlations were estimated by randomly shuffling the behavioral and cognitive data among participants (10,000 times). The FDR procedure was used to correct for multiple comparisons across all behavioral and cognitive domains at the level of *q* < 0.05 for the VBNs and DBNs, respectively. For each significant correlation, the contribution of a given edge was defined as its weight to form the PLS1. Furthermore, a BBS modeling method (33) was used to explore whether the morphological WM networks were predictive of individual performance in each behavioral and cognitive domain. First, principal component analysis was used for dimensionality reduction by retaining components that explained 80% variance in the interregional morphological similarities. Then, a linear regression model was fitted between the expression scores of the retained components and behavioral and cognitive data at each domain, and used to predict behavioral and cognitive outcomes for unseen participants. Finally, we calculated the Pearson correlation between actual scores and predicted values for each behavioral and cognitive domain. To assess the performance of the BBS model, a 10-fold cross-validation procedure was used. Since a single cross-validation might be sensitive to a particular split of the data into folds (85), the 10-fold cross-validation procedure was repeated 100 times, and the resulting mean Pearson correlation coefficient was reported for each behavioral and cognitive domain. To test whether the Pearson correlation coefficients were significantly higher than random operations, a nonparametric permutation testing procedure was performed by reshuffling the group labels of participants and repeating the 10-fold cross-validation procedure (10,000 times). The FDR procedure was used to correct for multiple comparisons across all behavioral and cognitive domains at the level of *q* < 0.05 for the DBNs and VBNs, respectively. For each significant correlation, the contribution of a given edge was calculated as the mean value of the product of coefficient in principal component analysis with beta value in linear regression model across all folds and repetitions.

### Heritability of morphological WM networks (HCP dataset, twin participants)

A genetic ACE model was used to investigate the extent to which morphological WM networks were genetically controlled. In a genetic ACE model, the variance of a phenotypic variable is assumed to be the sum of additive genetic contribution (A), common (C) and unique environment (E) contribution (86). Formally, narrow-sense heritability is defined as the proportion of phenotypic variance that is attributed to genetic factors:

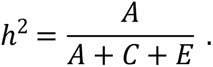

The heritability was estimated for each edge in the VBNs and DBNs with the APACE package (87). Significance level of each edge’s heritability was estimated by randomly shuffling the labels (MZ or DZ) of each pair of twins (10,000 times) to generate an empirical distribution. The FDR procedure was used to correct for multiple comparisons across all edges at the level of *q* < 0.05 for the VBNs and DBNs, respectively.

We further examined the relationships between the edge’s heritability and contribution to the observed significant phenotypic associations of the morphological WM networks via Spearman correlation. Significance levels of the correlations were estimated through simulating the heritability matrix via Moran spectral randomization to account for spatial autocorrelation (10,000 times) (75). The FDR procedure was used to correct for multiple correlations at the level of *q* < 0.05 for the VBNs and DBNs, respectively.

### Hamodynamic correlates of morphological WM networks (HCP dataset, unrelated participants)

#### Preprocessing of R-BOLD-fMRI images

For individual R-BOLD-fMRI images, the HCP fMRIVolume pipeline was used to generate minimally preprocessed 4D time series that included gradient distortion correction, motion correction, field map-based echo-planar imaging distortion correction, registration of functional data to structural scan, non-linear registration into the standard MNI space and grand-mean intensity normalization (88). After these steps, individual functional images further underwent band-pass filtering (0.01 to 0.08 Hz) and removal of nuisance covariates [24-parameter head motion profiles (89), cerebrospinal fluid signals and global signals], both of which were implemented in a single regression model to avoid reintroducing artifacts (90). The cerebrospinal fluid signals were calculated according to a prior probability map of cerebrospinal fluid as released in the SPM12 package (threshold = 0.9).

#### Construction of hamodynamic WM networks

The hamodynamic WM networks were constructed using the same WM parcellation atlas as the morphological WM networks. First, regional mean time series was extracted for each ROI by averaging signals of all voxels within the ROI for each volume. The resulting time series were then correlated with each other to generate a hamodynamic coherence matrix for each participant. Finally, the hamodynamic coherence matrices were averaged across participants to derive a group-level mean hamodynamic WM network.

#### Relationship between morphological and hamodynamic WM networks

We used the communication model, a multilinear regression model based on simple dynamical processes (34), to predict the group-level mean hamodynamic coherence profile, with group-level mean similarity profile and communicational organization (shortest path length and communicability) derived from the morphological WM networks as predictors (35). We constructed the model at whole brain level and for each node. Specifically, communicational organization of a node combined both centralized (shortest path length) and decentralized (communicability) policies requiring global and local knowledge of a network’s topology, respectively (34). Shortest path length was calculated as the minimum number of edges required to go from one node to another, and communicability was defined as the weighted sum of all paths and walks between two nodes (91). Shortest path length and communicability were both estimated from individual binary WM networks and averaged across participants. The *AR^2^* value was used to assess the performance of the multilinear regression model. Significance levels of the *AR^2^* values were estimated by predicting the hamodynamic coherence simulated via Moran spectral randomization (10,000 times) (75). The FDR procedure was used to correct for multiple comparisons across all nodes at the level of *q* < 0.05 for the DBN and VBN, respectively.

#### Association of the relationship between morphological and hamodynamic WM networks with functional hierarchy

To explore whether the relationship between morphological and hamodynamic WM networks follows functional hierarchies, we calculated hamodynamic coherence gradients of WM using the BrainSpace toolbox (92) in a similar way as for functional networks of gray matter (93–95). Briefly, an affinity matrix was first derived by calculating the cosine similarity between each pair of regional group-level mean hamodynamic coherence profiles containing the top 10% of correlations for each node. Then, diffusion map embedding, a nonlinear dimensionality reduction technique, was used to identify principle gradients that accounted for primary variations in the distribution of hamodynamic coherence across different regions. Finally, we calculated the Spearman rank correlation between the nodal *AR^2^* values derived above and the first hamodynamic coherence gradient of WM (Z-transformed). Significance level of the correlation was estimated through simulating the nodal *AR^2^* values via Moran spectral randomization to account for spatial autocorrelation (10,000 times) (75).

### Metabolic correlates of morphological WM networks **(MU dataset)**

#### Preprocessing of R-FDG-PET images

The R-FDG-PET images were preprocessed using the SPM12 package (https://www.fil.ion.ucl.ac.uk/spm/software/spm12/). First, we retained the 225 volumes commencing from the 30-minute time point, which matched the start of the R-BOLD-fMRI data acquisition in the dataset (29). Then, individual images were corrected for head motion, and normalized into the standard MNI space via transformation fields derived from tissue segmentation of structural images. One participant was excluded from further analyses due to the poor performance of PET-MRI co-registration. Finally, individual images were processed using a spatiotemporal gradient filter, the convolution of a 3-dimensional Gaussian filter in the spatial domain (standard deviation, one voxel) and a 1-dimensional Gaussian filter in the time domain (standard deviation, two frames), to estimate short-term changes in glucose uptake from the cumulative glucose uptake that was measured (29).

#### Construction of metabolic WM networks

The metabolic WM networks were constructed using the same WM parcellation atlas as the morphological WM networks. First, regional mean time series was extracted for each ROI by averaging signals of all voxels within the ROI for each volume. The resulting time series were then correlated with each other to generate a metabolic synchronization matrix for each participant. Finally, the metabolic synchronization matrices were averaged across participants to derive a group-level mean metabolic WM network.

#### Relationship between morphological and metabolic WM networks

The relationship between morphological and metabolic WM networks were examined using the multilinear regression model in the same manner as for the relationship between morphological and hamodynamic WM networks.

#### Association of the relationship between morphological and metabolic WM networks with regional metabolism

To test whether the relationship between morphological and metabolic WM networks was related to regional metabolism, we calculated the Spearman rank correlation between the nodal *AR^2^* values derived above and regional static metabolic values. For a given region, the static metabolic value was calculated as the mean metabolic activity across all time points and participants. Significance level of the correlation was estimated through simulating the nodal *AR^2^* values via Moran spectral randomization to account for spatial autocorrelation (10,000 times) (75).

### Chemoarchitectonic correlates of morphological WM networks (JuSpace dataset and HCP dataset, unrelated participants)

#### Construction of chemoarchitectonic WM network

Mounting evidence indicates the existence of multiple neurotransmitters in WM tracts, such as serotonergic, glutamatergic, dopaminergic, GABAergic, purinergic, adrenergic and cholinergic signaling (36). Thus, we constructed a chemoarchitectonic WM network to examine chemoarchitectonic correlates of the morphological WM networks. The chemoarchitectonic WM network was constructed using the same WM parcellation atlas as the morphological WM networks. First, the mean intensity was extracted for each ROI from each neurotransmitter receptor and transporter map in the JuSpace dataset. The resulting regional profiles were then Z-transformed to make comparability across different neurotransmitter receptors and transporters. Finally, the Z-transformed regional neurotransmitter receptor and transporter profiles were correlated with each other to generate a chemoarchitectonic covariance matrix.

#### Relationship between morphological and chemoarchitectonic WM networks

The relationship between morphological and chemoarchitectonic WM networks were examined using the multilinear regression model in the same manner as for the relationship between morphological and hamodynamic WM networks.

#### Association of the relationship between morphological and chemoarchitectonic WM networks with regional neurotransmitter intensity

To investigate whether the relationship between morphological and chemoarchitectonic WM networks was related to regional intensity of certain neurotransmitter receptor and transporter, we calculated the Spearman rank correlation between the nodal *AR^2^* values derived above and regional mean intensity of each neurotransmitter receptor and transporter. Significance levels of the correlations were estimated through simulating the nodal *AR^2^* values via Moran spectral randomization to account for spatial autocorrelation (10,000 times) (75). The FDR procedure was used to correct for multiple comparisons across all neurotransmitter receptors and transporters at the level of *q* < 0.05 for the DBN and VBN, respectively.

### Genetic correlates of morphological WM networks (AHBA dataset)

#### Preprocessing of gene data

Standardized workflows (96) were used to preprocess the gene data in the AHBA dataset with the abagen toolbox (version 0.1.3; https://github.com/rmarkello/abagen) (97). First, we updated the probe-to-gene annotations using up-to-date information. Then, intensity-based filtering was applied to exclude probes that did not exceed background noise in more than 50% of the samples. Afterwards, a representative probe was selected for each gene that had the most consistent pattern of regional variations across the six donor brains as quantified by a measure called Differential Stability (98, 99). To assign gene expression samples to the WM ROIs, we excluded samples further than 2 mm away from any voxel in the parcellation, and assigned each of the remaining samples to its nearest region according to the minimum distance between the sample and any voxel in a region. Finally, gene expression levels of the remaining samples were normalized for each donor by applying a scaled robust sigmoid normalization for every sample across genes and for every gene across samples in order to assess the relative expression of each gene across regions while controlling for donor-specific differences in gene expression.

#### Construction of transcriptional WM network

The transcriptional WM network was constructed using the same WM parcellation atlas as the morphological WM networks. First, the mean expression level of each gene was extracted for each ROI by averaging across samples of all donors assigned to the same region. The resulting regional gene expression profiles were then correlated with each other to generate a transcriptional gene co-expression network.

#### Relationship between morphological and transcriptional WM networks

The relationship between morphological and transcriptional WM networks were examined using the multilinear regression model in the same manner as for the relationship between morphological and hamodynamic WM networks.

#### Association of the relationship between morphological and transcriptional WM networks with regional gene expression

To investigate whether the relationship between morphological and transcriptional WM networks was related to regional expression of certain genes, we performed the PLS regression to predict the nodal *AR^2^* values derived above with regional expression levels of all genes. The PLS1 was the linear combination of regional expression levels of all genes that exhibited the strongest correlation with the nodal *AR^2^* values. Significance level of the correlation was estimated by re-running the PLS regression for nodal *AR^2^* values simulated via Moran spectral randomization to account for spatial autocorrelation (10,000 times) (75). If a significant correlation was observed, the weights of all genes to form the PLS1 were Z-transformed, and genes with an absolute Z-score > 1.64 were considered to strongly contribute to the relationship between the morphological and transcriptional WM networks.

#### Gene ontology enrichment analysis of genes contributing to the relationship between morphological and transcriptional WM networks

For the identified genes that strongly contributed to the relationships between the morphological and transcriptional WM networks, we performed GO enrichment analysis to search for their related GO terms. First, we downloaded the biological process-related GO term hierarchy files and annotation files for Homo sapiens (version April 17, 2019) from https://figshare.com/s/71fe1d9b2386ec05f421 (100). Then, we ran the gene-to-category annotations, processed the hierarchy relationships between GO terms, and restricted our analysis to the GO terms with 10 - 1,000 gene annotations (101, 102). To reduce the false-positive rate in GO enrichment analysis, a spatial ensemble null model was used (100). Specifically, for each of the resulting GO terms, an enrichment coefficient was calculated. To estimate the significance levels of the enrichment coefficients, we re-ran the PLS regression for nodal *AR^2^* values simulated via Moran spectral randomization (75), and re-calculated the enrichment coefficient for each GO term. These procedures were repeated 10,000 times to generate a null distribution for each GO term, based on which a *P*-value was calculated as the proportion of repetitions for which the resulting enrichment coefficient exceeded or equaled the real observation. Significant GO terms were determined after correcting for multiple comparisons with the FDR procedure at the level of *q* < 0.05. Notably, the GO enrichment analysis was performed for genes that positively and negatively contribute to the relationship between the morphological and transcriptional WM networks, respectively (i.e., PLS1+ and PLS1-genes).

#### Cell type-specific aggregation analysis of genes contributing to the relationship between morphological and transcriptional WM networks

For the identified genes that strongly contributed to the relationships between the morphological and transcriptional WM networks, we explored the effects of regional variations in cellular architecture on their contributions. First, we calculated an enrichment coefficient for each of the seven canonical cell classes: excitatory neurons, inhibitory neurons, oligodendrocyte progenitor cells, astrocytes, endothelial cells, microglia and oligodendrocytes (103). The significance levels of the enrichment coefficients were then estimated in a similar manner as for the GO enrichment analysis. Finally, the FDR procedure was used to correct for multiple comparisons across all cell classes at the level of *q* < 0.05. Again, the cell type-specific aggregation analysis was separately performed for the PLS1+ and PLS1-genes.

### Clinical correlates of morphological WM networks (MS and NMOSD multicentric dataset)

Clinical correlates of morphological WM networks were explored by examining disease-related alterations in interregional morphological similarity, associations of altered morphological similarity with clinical variables, and classification and differentiation of MS and NMOSD. Before these analyses, morphological WM networks were harmonized for site effects using the toolbox of ComBat (https://github.com/Jfortin1/ComBatHarmonization/tree/master/Matlab).

#### Disease-related alterations in interregional morphological similarity

The threshold-free network-based statistics (TFNBS) approach (104) was used to examine the differences in interregional morphological similarity among the three groups. Specifically, one-way ANCOVA was first performed for each edge in the morphological WM networks with group as a between-subject factor and sex and age as covariates of non-interest. This resulted in a 48 × 48 *F* statistic matrix, which was then enhanced with the TFNBS approach by combining the threshold-free cluster enhancement and network-based statistic (105). The extension and height enhancement parameters were set to 0.5 and 2, respectively, based on the recommendations from Baggio and colleagues (104). Significance levels of the enhanced *F* statistics were estimated through a nonparametric permutation testing procedure (10,000 times), and corrected for multiple comparisons by comparing each edge’s statistic to the distribution of the maximum statistic of all edges under the null hypothesis. For edges showing significant group effects, post hoc tests were further performed via the TFNBS approach with independent two-sample *T* test.

#### Associations of altered interregional morphological similarity with clinical variables

Spearman rank correlation was used to examine the relationships between the altered interregional morphological similarity and clinical variables (disease duration and Expanded Disability Status Scale score) in each patient group. Effects of age and sex were removed from the altered interregional morphological similarity via a linear regression model. The FDR procedure was used to correct for multiple comparisons in each patient group at the level of *q* < 0.05.

#### Classification and differentiation of MS and NMOSD

To examine the potential of morphological WM networks in diagnosing and differentiating the two diseases, we trained a binary linear support vector machine classifier for each pair of groups with all interregional morphological similarities as original features. Before constructing the classifiers, principal component analysis was used for dimensionality reduction by retaining components that explained 80% variance in the original features. To assess the performance of the classifiers, a 10-fold cross-validation was used. Since a single cross-validation might be sensitive to a particular split of the data into folds (85), the 10-fold cross-validation procedure was repeated 100 times, and the resulting mean accuracy was reported for the classification between each pair of groups. To test whether the classification accuracies were significantly higher than random operations, a nonparametric permutation testing procedure was performed by reshuffling the group labels of participants and repeating the 10-fold cross-validation procedure (10,000 times).

## Acknowledgements

This work was supported by the National Natural Science Foundation of China (No. 81922036), Key Realm R&D Program of Guangdong Province (No. 2019B030335001), Key Realm R&D Program of Guangzhou (No. 202007030005) and National Social Science Foundation of China (No. 20&ZD296).

## Data availability statement

All data that support the findings of this study are from publicly available datasets (HCP dataset: www.humanconnectome.org; HNU dataset: http://dx.doi.org/10.15387/fcp_indi.corr.hnu1; SWU dataset: http://dx.doi.org/10.15387/fcp_indi.retro.slim; MU dataset: https://openneuro.org/datasets/ds002898/versions/1.1.0; JuSpace datset: https://github.com/juryxy/JuSpace; AHBA dataset: http://human.brain-map.org/) except for the MS and NMOSD multicentric dataset, which are available from the corresponding author upon reasonable request. Due to the nature of this research, participants of the MS and NMOSD multicentric dataset did not agree for their data to be shared publicly.

## Code availability

The neuroimaging preprocessing software is freely available (CAT12, http://www.neuro.uni-jena.de/cat/; SPM12, https://www.fil.ion.ucl.ac.uk/spm/software/spm12). Code for all the analyses excluding the neuroimaging preprocessing is available at https://github.com/Junle-1995/Morphological-WMNs. Related toolboxes include GRETNA (https://www.nitrc.org/projects/gretna; topological analysis), MRM (http://www.click2go.umip.com/i/software/mrm.html; two-way repeated ANOVA), APACE (http://warwick.ac.uk/tenichols/apace; heritability analysis), Brainspace (https://brainspace.readthedocs.io/en/latest/index.html; Moran spectral randomization and gradient calculation), abagen (version 0.1.3; https://github.com/rmarkello/abagen; preprocess for gene data), GeneCategoryEnrichmentAnalysis (https://github.com/benfulcher/GeneCategoryEnrichmentAnalysis; gene ontology enrichment analysis).

## Author contributions

JW designed the study; JL, ZL, YY and ZL analyzed data; JL wrote the manuscript; JW and YL revised the manuscript.

